# A CRISPR-Nonhomologous End-Joining-based strategy for rapid and efficient gene disruption in *Mycobacterium abscessus*

**DOI:** 10.1101/2024.01.29.577284

**Authors:** Sanshan Zeng, Yanan Ju, Md Shah Alam, Ziwen Lu, H.M. Adnan Hameed, Lijie Li, Xirong Tian, Cuiting Fang, Xiange Fang, Jie Ding, Xinyue Wang, Jinxing Hu, Shuai Wang, Tianyu Zhang

## Abstract

*Mycobacterium abscessus*, a fast-growing, non-tuberculous mycobacterium resistant to most antimicrobial drugs, causes many types of serious infections in humans, posing a significant public health challenge. Currently, effective genetic manipulation tools for *M. abscessus* are still being developed, which hampers research and therapeutic development. However, the clustered regularly interspaced short palindromic repeats (CRISPR) - associated protein (Cas) systems have emerged as promising methods for generating highly specific double-strand breaks (DSBs) in its genome. These DSBs can be repaired by the error-prone nonhomologous end joining (NHEJ) mechanism, facilitating targeted gene editing. Here, our study marks a pioneering application of the CRISPR-NHEJ strategy in *M. abscessus*. Additionally, we discovered that NrgA from *Mycobacterium marinum* is crucial for the repair of DSBs caused by the CRISPR-Cas system in *M. abscessus*. Finally, contrary to previous findings, our study also indicates that inhibiting or overexpressing homologous recombination/single-strand annealing significantly decreases the efficiency of NHEJ repair in *M. abscessus*. This discovery challenges established perspectives and suggests that the NHEJ repair in *M. abscessus* may require the involvement of components from homologous recombination and single-strand annealing, demonstrating the complex interactions among the three DSB repair pathways in *M. abscessus*.

**Impact statement:** There are still very few genetic manipulation tools available for *Mycobacterium abscessus*. Here we report the successful application of CRISPR-Cas12a-assisted nonhomologous end joining (NHEJ) in efficient gene editing in *M. abscessus*. Contrary to previous research suggesting that homologous recombination (HR) inhibition may enhance such editing efficiency in other Mycobacterium species, our results showed that disruption or overexpression of either HR or single-strand annealing not only failed to enhance but also significantly reduced the gene editing efficiency in *M. abscessus*. This suggests that NHEJ repair in *M. abscessus* may require components from both HR and single-strand annealing, highlighting a complex interaction among the DSB repair pathways in *M. abscessus*.

## INTRODUCTION

The prevalence of lung infections caused by non-tuberculous mycobacteria (NTM) has been rising annually, posing an escalating threat to public health. Among these NTM, *M. abscessus* is one of the primary causative agents in both the United States and the humid regions of southern coastal China^1–3^. Besides contagious pulmonary infections, *M. abscessus* can infect the skin, soft tissues, bone and various other parts of the human body^4^. The intrinsic resistance of *M. abscessus* to a broad spectrum of antibiotics not only complicates the treatment, but often requires the use of multidrug regimens and prolonged therapies^5^. The threat posed by *M. abscessus* necessitates the urgent development of genetic tools for drug target identification, resistance mechanism elucidation, and vaccine research.

The Clustered Regularly Interspaced Short Palindromic Repeats (CRISPR) – CRISPR-associated protein (Cas) systems have been widely applied as efficient gene editing tools in various organisms^6^. Among these systems, CRISPR-associated protein 9 (Cas9) and Cas12a (formerly known as Cpf1) are the most common and important nucleases^7, 8^. Guided by the corresponding CRISPR-RNA, Cas9 and Cpf1 can precisely recognize and bind to the target DNA sequence. This process is dependent on the complementary base pairing between the CRISPR-RNA and the target DNA sequence. Once matched, the CRISPR effector nuclease induces double-strand breaks (DSBs) at designated loci within the target DNA. However, DSBs are lethal to organisms because they can severely compromise genomic integrity and stability^9, 10^.

To address this damage, organisms have evolved various mechanisms to repair DSBs^11^. In contrast to other prokaryotes, the DSB repair system in mycobacteria displays a high degree of evolutionary complexity and diversity, similar to that of *Saccharomyces cerevisiae*, and includes redundant repair components^12^. Therefore, mycobacteria can repair DSBs through multiple distinct mechanisms. Among these mechanisms, homologous recombination (HR) is the most accurate DSB repair process. However, it requires an additional intact homologous template. Single-strand annealing (SSA), another DSB repair mechanism, primarily occurs in the presence of direct repeat sequences flanking the DSBs^13^. Mycobacteria also possess the non-homologous end joining (NHEJ) repair mechanism. Unlike the complex NHEJ system in eukaryotic cells, mycobacterial NHEJ consists of only two primary components: the end-binding and end-bridging DNA binding protein Ku and the multifunctional enzyme LigD^14, 15^. Firstly, the Ku protein binds to the DNA termini, then LigD facilitates minimal end processing and directly ligates the processed termini without requiring a homologous template, making this type of repair error-prone. Through HR, SSA, or NHEJ repair mechanisms, mycobacteria effectively repair DSBs^10^. The application of these repair mechanisms, as well as the integration of CRISPR-Cas systems with these mechanisms for gene editing, is gaining popularity in mycobacteria^16–20^.

Methods for genetic manipulation of Mycobacteria are constantly being improved^21^. Initially, genetic manipulations such as the expression of *gp60* and *gp61* from the mycobacteriophage Che9c have been accomplished for enhanced recombination in *Mycobacterium smegmatis, Mycobacterium tuberculosis*^16^, and *Mycobacterium abscessus*^20^. CRISPR-assisted recombineering has further enabled the effectual construction of marker-free recombinants in mycobacteria^17^. After successful implication in *M. smegmatis*, this technique has recently been applied successfully for the first time by us in *M. abscessus*^19^. Additionally, the template-independent and error-prone DSB repair mechanism NHEJ, presents the potential for genetic manipulation together with CRISPR. To date, the CRISPR-NHEJ gene manipulation approach has been proven effective in *M. smegmatis* and *M. tuberculosis,* however, no study reported the application of these strategies on *M. abscessus* so far^18^.

The *nrgA*(NHEJ-related gene A), locates between the *ku* and *ligD* genes in the *M. marinum* chromosome, belonging to the phosphofructokinase B-type (PfkB) family of sugar kinase encoding genes (*MMAR_4574*). As a phosphofructokinase, the structure of NrgA has recently been elucidated^22^. It uses phosphorylated sugars as substrates and catalyzes the conversion of fructose-6-phosphate to fructose-1,6-bisphosphate. However, this protein has no homolog in *M. abscessus*. Research indicates that in *M. marinum* and *M. tuberculosis*, mutations in *nrgA* significantly reduce the efficiency of CRISPR-NHEJ-based gene editing^18^. This suggests that *nrgA* may play an important role in the NHEJ repair pathway in *M. marinum*. However, the specific mechanisms remain unclear.

To achieve efficient gene knockout using a CRISPR-NHEJ strategy in *M. smegmatis* and *M. tuberculosis*, it was necessary to regulate their inherent HR systems^18^. This can be accomplished by expressing the negative regulator of *recA*, *recX*, or by directly knocking out the key HR pathway gene, *recA*, to suppress or eliminate HR^23, 24^. Under such conditions, a significant increase in gene editing efficiency is observed. Conversely, when the HR repair is not regulated, the efficiency of gene editing using the CRISPR-NHEJ strategy is extremely low. Based on these findings, researchers believed that HR and NHEJ are competitive pathways in the repair of DSBs induced by the CRISPR-Cas system in mycobacteria^18^.

In the first part of this study, we report for the first time that gene editing using the CRISPR-NHEJ strategy is efficient in *M. abscessus*. Then, we discovered that NrgA, a component of the NHEJ system in *M. marinum*, is indispensable for the NHEJ-mediated repair of DSBs induced by the CRISPR-Cas system in *M abscessus*. In the final part of our study, based on previous research, we investigated the impact of *recX* on the gene editing efficiency of CRISPR-NHEJ strategy in *M. abscessus* and *M. smegmatis* and discovered differing outcomes. We further explored the relationship among the three DSB repair mechanisms in *M. abscessus*. Our findings indicate a complex interplay among these three repair mechanisms during the DSB repair process, laying the foundation for future studies to further elucidate their relationships.

## RESULTS

### Expression of *M. marinum* NHEJ components enabled effective gene editing at CRISPR-Cpf1-induced double-strand breaks and improved the survival rate in *M. abscessus*

Compared to the complex NHEJ system in eukaryotes, *M. marinum* possesses a relatively simple NHEJ system, consisting of only LigD, Ku, and NrgA. It has been demonstrated that the endogenous NHEJ system in *M. marinum* is capable of effectively repairing DSBs induced by the CRISPR-Cas system, suggesting that it is a potent NHEJ repair^18^. Therefore, we first constructed a plasmid, pNHEJ-Cpf1, which simultaneously expresses CRISPR-Cpf1 and the complete NHEJ repair system (Ku, NrgA, LigD) from *M. marinum* (Supplementary Figure S1). In this plasmid, the expression of Cpf1 is induced by anhydrotetracycline, while the NHEJ components from *M. marinum* are under the control of their native promoters.

To investigate the effect of the *M. marinum* NHEJ system expression on *M. abscessus* gene editing efficiency, we transformed *M. abscessus* cells with plasmids pNHEJ-CPF1 or pJV53-Cpf1, respectively. In the absence of acetamide induction, pJV53-Cpf1 only expresses Cpf1 in *M. abscessus* cells (plasmids used in this study are listed in Supplementary Table S1). After obtaining *M. abscessus* cells carrying the respective plasmids, we used another plasmid, pCR-ZEO-3513c, to express the crRNA targeting the *MAB_3513c* gene. We then used this dual-plasmid system to attempt the knockout of the *Mab_3513c* gene and observed the differences in survival rate and gene editing efficiency (Fig. 1A). After transforming the pCR-ZEO-3513c plasmid, the cultures were plated on 7H11 agar plates containing the appropriate antibiotics and 200 ng/ml ATc, with plates lacking ATc serving as the control. The survival rate was defined as the ratio of CFUs on plates with ATc to those on plates without ATc. Compared to the survival rate of the pJV53-Cpf1 group, which is only 47.5 ± 3.54%, the survival rate of the colonies in the pNHEJ-Cpf1 group is significantly higher at 71 ± 2.83% (*P* = 0.0156, Fig. 1B). This suggests that the expression of the *M. marinum* NHEJ system repaired some of the DSBs caused by CRISPR-Cpf1, thereby restoring a portion of the colonies. We further analyzed the gene editing efficiency (editing efficiency was calculated by measuring the proportion of colonies with mutations at the target site among the total randomly selected colonies) of *M. abscessus* obtained through the CRISPR-NHEJ strategy. First, we randomly selected several single colonies from the ATc-containing plates and used the primer pair Mab_3513c-F/R to amplifies the *MAB_3513c* gene of the individual colonies (Fig. 1C). The electrophoresis results, as shown in Figure 1D, reveal that some single colonies exhibit PCR amplification products significantly smaller than those of the wild-type strain, or no amplification bands at all. These outcomes can be attributed to large DNA deletions that occurred during the NHEJ repair process.

**FIG 1.**
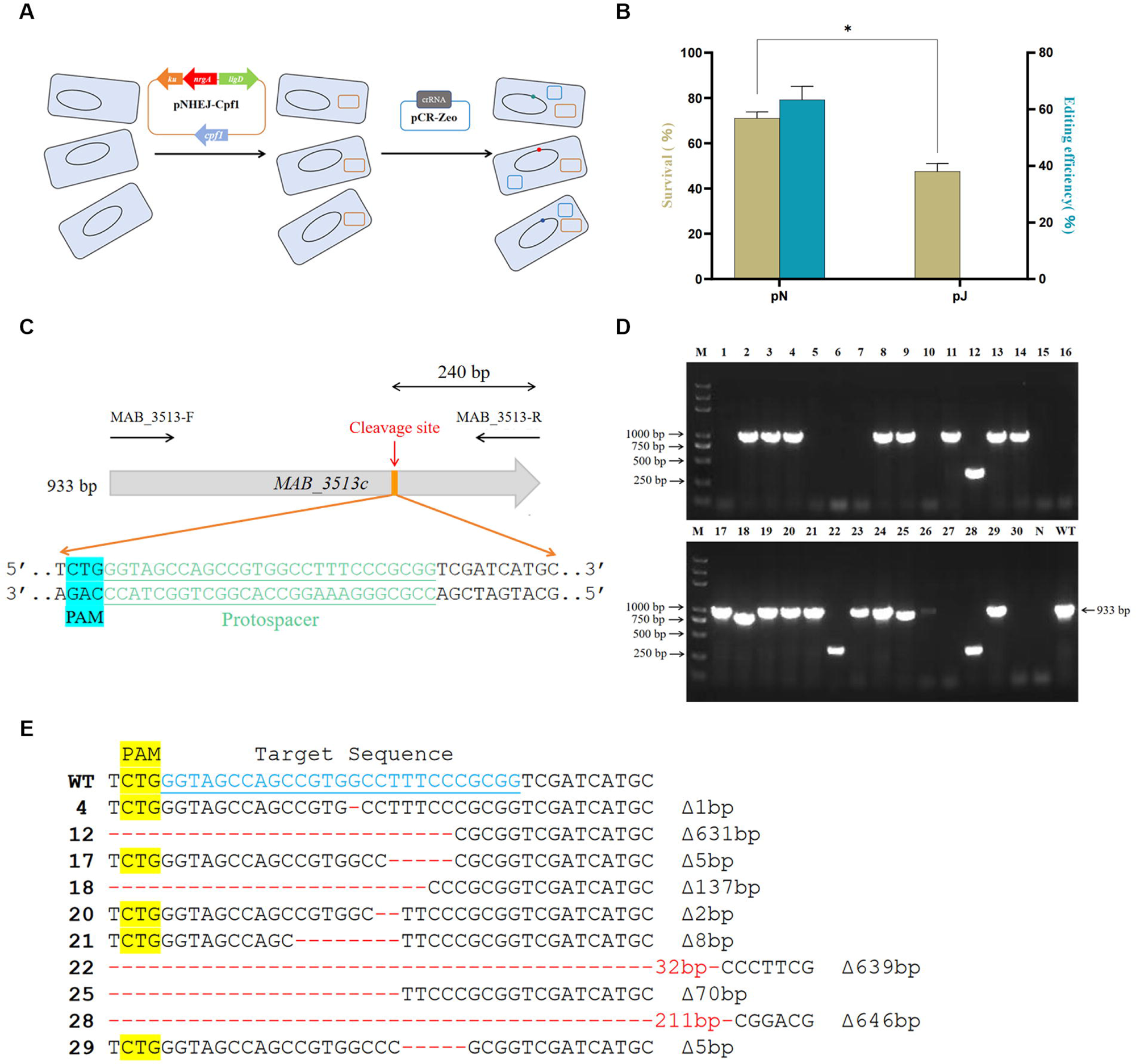
CRISPR-Cpf1-NHEJ-assisted gene editing in *M. abscessus.* (A) Schematic of CRISPR-Cpf1 coupled with the *M. marinum* NHEJ repair in *M. abscessus*. In the first step, Cpf1 and *M. marinum* NHEJ components were expressed in *M. abscessus.* Then, the crRNA-expressing plasmid pCR-Zeo was transformed into the cell. (B) The schematic diagram of the coding sequence, identification primers, cleavage sites, PAM, and protospacer sequence for *MAB_3513c*. (C) The Colony PCR results (agarose gel electrophoresis) of randomly selected clones for *MAB_3513c* deletion under the action of *M. marinum* repair system. Randomly select 30 single clones and amplify the *MAB_3513c* using the MAB_3513-F/R primer pair from Fig.1B. “N”, the PCR result for the negative control Mab^□3513c^, “P”, PCR result for the positive control of the *M. abscessus* wild-type strain. The expected amplicon length is 933 bp. (D) Expression of *M. marinum* NHEJ enables effective gene editing in *M. abscessus*. The plasmid expressing crRNA targeting *MAB_3513c* was introduced into *M. abscessus* strains harboring pNHEJ-Cpf1 or pJV53-Cpf1, respectively. Editing efficiency was calculated by measuring the proportion of colonies with mutations at the target site among the total randomly selected clones and survival rate was defined as the ratio of the number of CFUs obtained from plates with ATc to the number of CFUs obtained from plates without ATc. *, *P* < 0.05. (E) Sequencing results for the *MAB_3513c* target site in selected single colonies. Sequences highlighted with a yellow background indicate the PAM sequences, blue sequences represent the crRNA target sequences, red segments indicate single-base deletions, and the numbers correspond to the respective mutant strains. “WT” denotes the *M. abscessus* wild-type strain.

However, some single colonies produced PCR results nearly identical to those of the wild-type strain. This does not necessarily mean that the target sites were unchanged, as small changes of just a few base pairs cannot be differenate easily by electrophoresis alone. Therefore, we proceeded with sequencing these single colonies. Sequencing results for altered target sites are shown in Figure 1E. In addition to the large fragment deletions discernible from the PCR results, there were still many single colonies, such as colony 4、17、20、21、29, that exhibited only a few base pairs of DNA deletions. These DNA fragment deletions that are not multiples of three will cause frameshift mutations, ultimately leading to partial or complete inactivation of the gene. Statistical analysis revealed that the gene editing efficiency for knocking out the *MAB_3513c* gene using the CRISPR-NHEJ strategy reached 63.35 ± 4.74%. In contrast, none of the single colonies randomly selected from the pJV53-Cpf1 group showed any evidence of gene editing (Fig. 1D). This indicates that the endogenous NHEJ repair system in *M. abscessus* is not effective at repairing the DSBs caused by CRISPR-Cpf1. However, when the exogenous *M. marinum* NHEJ system was expressed, it facilitated error-prone repair of the DSBs induced by CRISPR-Cpf1.

After successfully knocking out the *MAB_3513c* gene using the CRISPR-NHEJ strategy, we aimed to assess the broader applicability of this method by targeting additional genes in *M. abscessus*. To date, we have successfully knocked out more than 40 genes in *M. abscessus* (Partially listed in Table 1). Depending on the specific gene being edited, the overall gene editing efficiency ranged from 40% to 90%, depending on the specific gene target, which is adequate for standard gene editing experiments. Compared to existing methods, this approach has significantly improved the efficiency of generating *M. abscessus* mutants.

### NrgA: a crucial factor in repairing CRISPR-Cas-induced double-strand breaks via NHEJ in *M. abscessus*

While *M. abscessus* has its own endogenous NHEJ repair system, our findings indicate that this system is ineffective at repairing DSBs generated by the CRISPR-Cpf1 system. Effective DSB repair in *M. abscessus* requires the introduction of the exogenous NHEJ repair system from *M. marinum*.

Compared to *M. abscessus*’s NHEJ repair system, the *M. marinum* NHEJ system includes not only LigD and Ku but also NrgA (Fig 2A). In *M. marinum*, the presence of NrgA has been shown to improve the editing efficiency of the CRISPR-NHEJ strategy^18^. Therefore, two factors may necessitate the introduction of the *M. marinum* NHEJ system for CRISPR-NHEJ gene editing: differences in LigD and Ku between *M. abscessus* and *M. marinum*, and the presence of the additional NrgA factor in *M. marinum*. To address this issue, we first constructed plasmids containing only part of the *M. marinum* NHEJ components, namely, *ku*, *nrgA*, *ligD*, *ku+nrgA*, *nrgA+ligD*, or *ligD+ku* from *M. marinum*, along with Cpf1. These correspond to the plasmids pK-Cpf1, pA-Cpf1, pD-Cpf1, pAK-Cpf1, pDA-Cpf1, and pDK-Cpf1, respectively. We then observed the survival rates and gene editing efficiencies in different *M. abscessus* strains carrying these plasmids after CRISPR-NHEJ gene editing. Compared to the pNHEJ-Cpf1 group, the survival rates of these *M. abscessus* strains containing only part of the *M. marinum* NHEJ components all decreased, indicating a reduction in the number of single colonies recovered by NHEJ repair (Fig 2B). Further analysis of the gene editing efficiencies in these strains showed that in the group simultaneously introduced with *M. marinum ku* and *ligD*, we observed a significant decrease in gene editing efficiency compared to the pNHEJ-Cpf1 group, with only 15 ± 7.07% (*P* = 0.0151). In the groups introduced with either *M. marinum ku* or *ligD* alone, we did not find any single colonies that underwent gene editing. However, in the group expressing only *M. marinum nrgA*, we found that approximately 7.5 ± 3.54% of the single colonies underwent gene editing. This suggests that the endogenous NHEJ system of *M. abscessus* may perform low-efficiency repair of DSBs caused by the CRISPR-Cpf1 system in the presence of NrgA. In the groups co-expressing *M. marinum ku* and *nrgA* or co-expressing *M. marinum ligD* and *nrgA*, the gene editing efficiencies increased compared to the NrgA group, with efficiencies of 15 ± 7.07% and 22.5 ± 3.536%, respectively, indicating that in the presence of NrgA, exogenous *ku* and *ligD* may also participate in part of the endogenous NHEJ repair process (Fig 2B).

**FIG 2.**
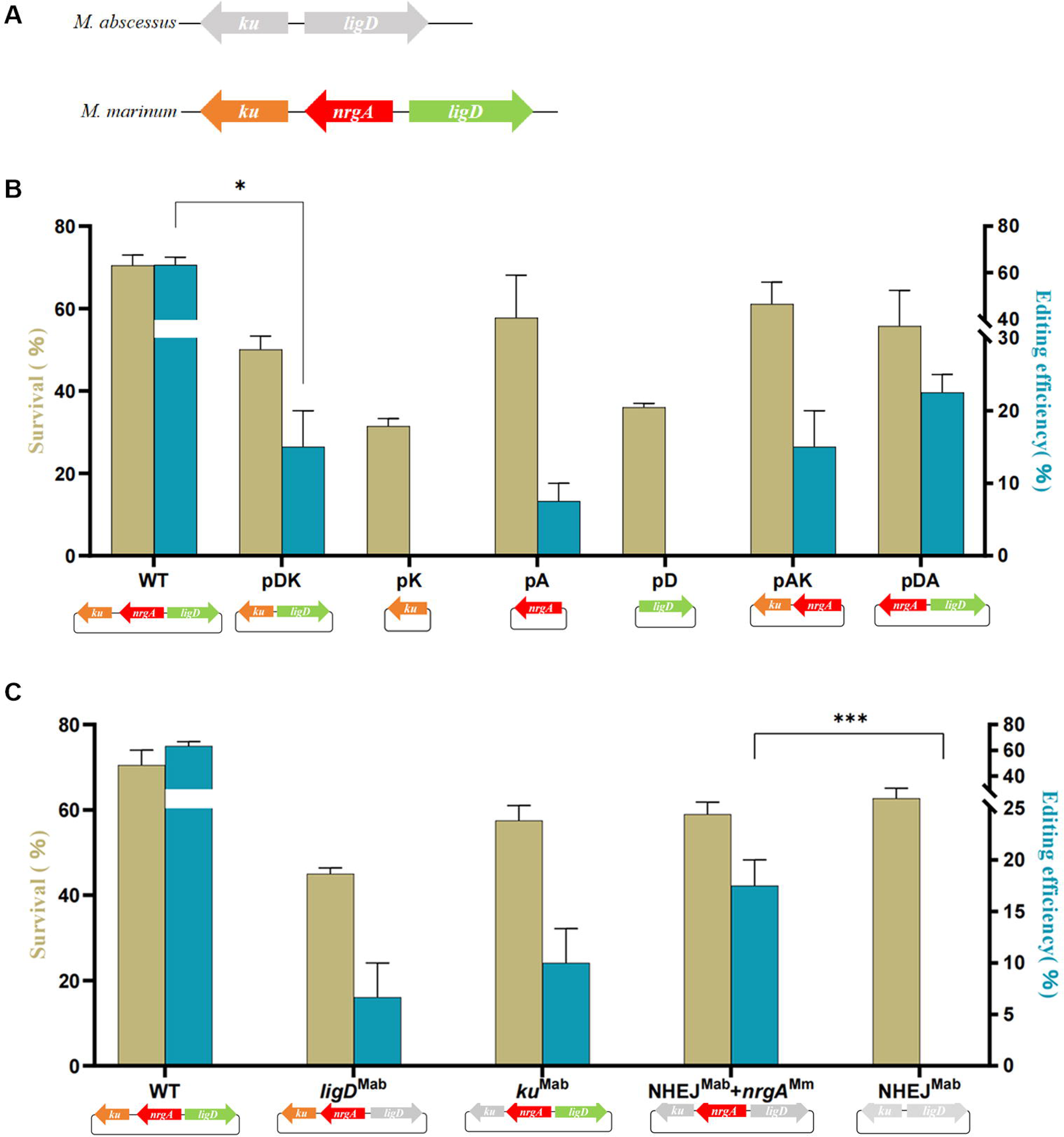
(A) Schematic of NHEJ components in *M. abscessus* and *M. smegmatis.* (B) CRISPR-Cas12a-NHEJ gene editing in *M. abscessus* with certain *M. marinum* NHEJ components. (C) CRISPR-Cas12a-NHEJ gene editing in *M. abscessus* with hybrid NHEJ components. Plasmids carrying the crRNA were electroporated into *M. abscessus* cells containing plasmids expressing different NHEJ components. Editing efficiency was calculated by measuring the proportion of colonies with mutations at the target site among the total randomly selected clones and survival rate was defined as the ratio of the number of CFUs obtained from plates with ATc to the number of CFUs obtained from plates without ATc. *, *P* < 0.05. ***, *P* < 0.001.

Additionally, we further explored the differences in survival rates and gene editing efficiencies by replacing the *M. marinum* NHEJ components with *M. abscessus* NHEJ components. Similarly, we constructed plasmids containing mixtures of *M. abscessus* and *M. marinum* NHEJ components: pNHEJ-Cpf1 (*ligD*^Mab^), pNHEJ-Cpf1 (*ku*^Mab^), pNHEJ-Cpf1 (NHEJ^Mab^ + *nrgA*^Mmr^), and a plasmid containing only *M. abscessus* NHEJ components without *M. marinum nrgA*, pNHEJ-Cpf1 (NHEJ^Mab^). We then conducted gene editing experiments in *M. abscessus* strains carrying these plasmids. The survival rates and gene editing efficiencies in groups transferred with other plasmids were all lower than those in the group transferred with the pNHEJ-Cpf1 plasmid (Fig 2C). Notably, the gene editing efficiency in the group where both *ligD* and *ku* of *M. marinum* were replaced with *ligD* and *ku* of *M. abscessus* (17.5 ± 3.536%) was higher than in the group where only *ligD* of *M. marinum* was replaced with *ligD* of *M. abscessus* (6.65 ± 4.716%) or where only *ku* of *M. marinum* was replaced with *ku* of *M. abscessus* (10 ± 4.709%). These results suggest that achieving high gene knockout efficiency requires that both LigD and Ku come from the same bacterial species. This also explains the slightly reduced gene editing efficiency observed in *M. abscessus* strains expressing mixed NHEJ components compared to those expressing only components from *M. marinum*. The increase in the proportion of mixed NHEJ components leads to a decrease in NHEJ repair efficiency (Fig 2B and C). Compared to the NHEJ^Mab^ + *nrgA*^Mmr^ group, we did not observe any single colonies undergoing targeted gene editing in the NHEJ^Mab^ group (*P* < 0.001), further confirming that *nrgA*^Mmr^, which is only present in *M. marinum*, plays an important role in the NHEJ repair process in *M. abscessus*.

### Inhibition of HR has different effects on the CRISPR-NHEJ gene editing efficiencies in *M. smegmatis* and *M. abscessus*

Previous studies have shown that inhibiting HR in *M. smegmatis* significantly enhances the efficiency of CRISPR-NHEJ editing, suggesting that HR and NHEJ might represent a competitive relationship during the DSB repair process in *M. smegmatis*. However, without additional expression of *recX*, we have already achieved efficient gene editing in *M. abscessus* using the CRISPR-NHEJ strategy. The differences between the reported results in *M. smegmatis* and our findings in *M. abscessus* prompted us to conduct further research. We first explored the impact of *recX* on the CRISPR-NHEJ gene editing efficiency in *M. smegmatis*. Here, we also investigated the differences in NHEJ repair of DSBs caused by different CRISPR effector proteins (Cas9, Cpf1) under the influence of *recX* in *M. smegmatis*. When using Cas9 as the effector protein, the gene editing efficiency in the group without *recX* expression was 49.45 ± 14.92%, while the gene editing efficiency with *recX* expression was 60 ± 14.14%, showing no significant difference (*P* = 0.5434, Fig. 3). In the case of using Cpf1 as the effector protein, the gene editing efficiency in the group without *recX* expression was 78.35 ± 11.81%, and with *recX* expression, it was 85 ± 7.071%, similarly showing no significant difference (*P* = 0.5650, Fig. 3). We found that whether using Cpf1 or Cas9 as the effector protein, the gene editing efficiency with or without *recX* expression showed no significant difference, and the gene editing efficiencies were all above 40%. This indicates that in *M. smegmatis*, we can achieve high-efficiency gene editing without the need for additional *recX* expression or manipulation of the host’s HR system, which is significantly different from previously reported results^18^. In *M. abscessus*, our preliminary research also indicated that achieving high-efficiency gene editing based on the CRISPR-NHEJ strategy does not require additional *recX* expression. We further observed the impact of overexpressing *recX* on the gene editing efficiency of the CRISPR-Cpf1-NHEJ strategy in *M. abscessus*. Surprisingly, after overexpressing *recX*, the gene editing efficiency significantly decreased to just 16.55 ± 2.192%, approximately 1/4 of the original efficiency. This precisely indicates that during the DSB repair process in *M. abscessus*, the NHEJ repair process may require the involvement of HR.

**FIG 3.**
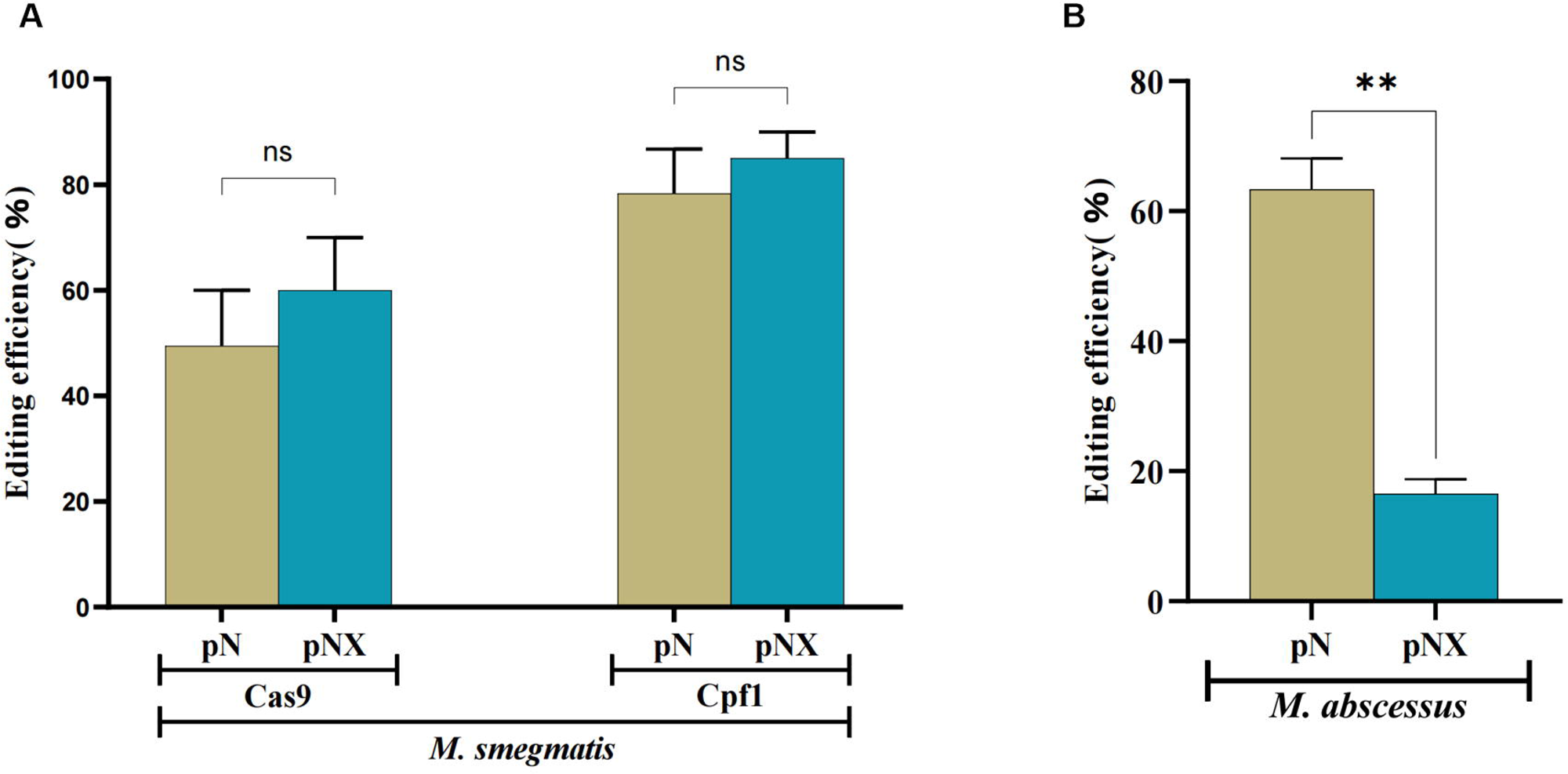
The effects of overexpressing the *recX* gene on the editing efficiency of *M. abscessus* and *M. smegmatis*. (A) The effects of overexpressing the *recX^Msm^* gene on the editing efficiency of *M. smegmatis*. The plasmid expressing sgRNA and Cas9 targeting *MSMEG_1946* was introduced into *M. smegmatis* strains harboring pNHEJ-SacB or pNHEJ-SacB(*recX*^Msm^), the plasmid expressing crRNA targeting *MSMEG_1946* was introduced into *M. smegmatis* strains harboring pNHEJ-Cpf1 or pNHEJX-Cpf1(*recX*^Msm^), respectively. (B) The effects of overexpressing the *recX^Mab^* gene on the CRISPR-Cas12a-NHEJ editing efficiency of *M. abscessus*. pN: plasmid with no *recX* but includes the sequence for the corresponding CRISPR effector proteins. pNX: plasmid with *recX* and includes the sequence for the corresponding CRISPR effector proteins. Editing efficiency was calculated by measuring the proportion of colonies with mutations at the target site among the total randomly selected clones and survival rate was defined as the ratio of the number of CFUs obtained from plates with ATc to the number of CFUs obtained from plates without ATc. ns, not significant. **, *P* < 0.01.

### In the DSB repair process of *M. abscessus*, NHEJ, HR, and SSA are in dynamic balance

In mycobacteria, the DSB repair mechanisms include not only HR and NHEJ but also SSA. While there have been studies on the impact of SSA on HR efficiency in mycobacteria^25^, the potential influence of SSA on NHEJ has been overlooked. Therefore, we simultaneously examined if disrupting HR or SSA would impact the NHEJ-dependent gene editing efficiency in *M. abscessus*. We constructed selection-marker-free, in-frame deletion knock-out strains of *recA*, *ruvB*, *recC*, *recO, ligD or ku* gene in *M. abscessus* using the CRISPR-assisted recombineering method as described in our previous study^19^. RecA plays a crucial role in HR and is thought to function at the initiation stage of HR repair, whereas RuvB promotes the branch migration of Holliday junctions to ensure accurate repair of DNA double-strand breaks^26–28^. The RecBCD complex, which includes RecC, is essential for SSA and is independent of RecA-mediated HR^13^. In addition, it was shown that the *recO* gene may play a role in both the HR and SSA pathways^29,30^. LigD and Ku are two crucial components of the endogenous NHEJ repair system in *M. abscessus*. Next, in these endogenous HR, SSA or NHEJ repair-deficient strains, we repeated the gene editing experiments performed in the wild-type strain as described above to observe the potential impact of endogenous HR, SSA or NHEJ repair defects on the gene editing efficiency.

Our results showed that the gene editing efficiencies were 3.33 ± 2.877%, 22.5 ±3.536%, 17.5 ± 3.536%, 7.5 ± 3.536% in the *recA*, ruvB, *recC* or *recO* knock-out strains, respectively (Fig. 4A). As expected, the gene editing efficiency in the *recA* knock-out strain was lower relative to the wild-type strain (*P* = 0.0004). Similarly, the gene editing efficiencies in the *ruvB*, *recC* or *recO* gene knock-out strains were significantly lower than that in the wild-type strain (*P* = 0.0103, *P* = 0.0082 and *P* = 0.0056, respectively, Fig. 1C). All these results supported that disruption of the HR or SSA repairing pathway would decrease the CRISPR-Cas12a-assisted NHEJ gene editing efficiency.

**FIG 4.**
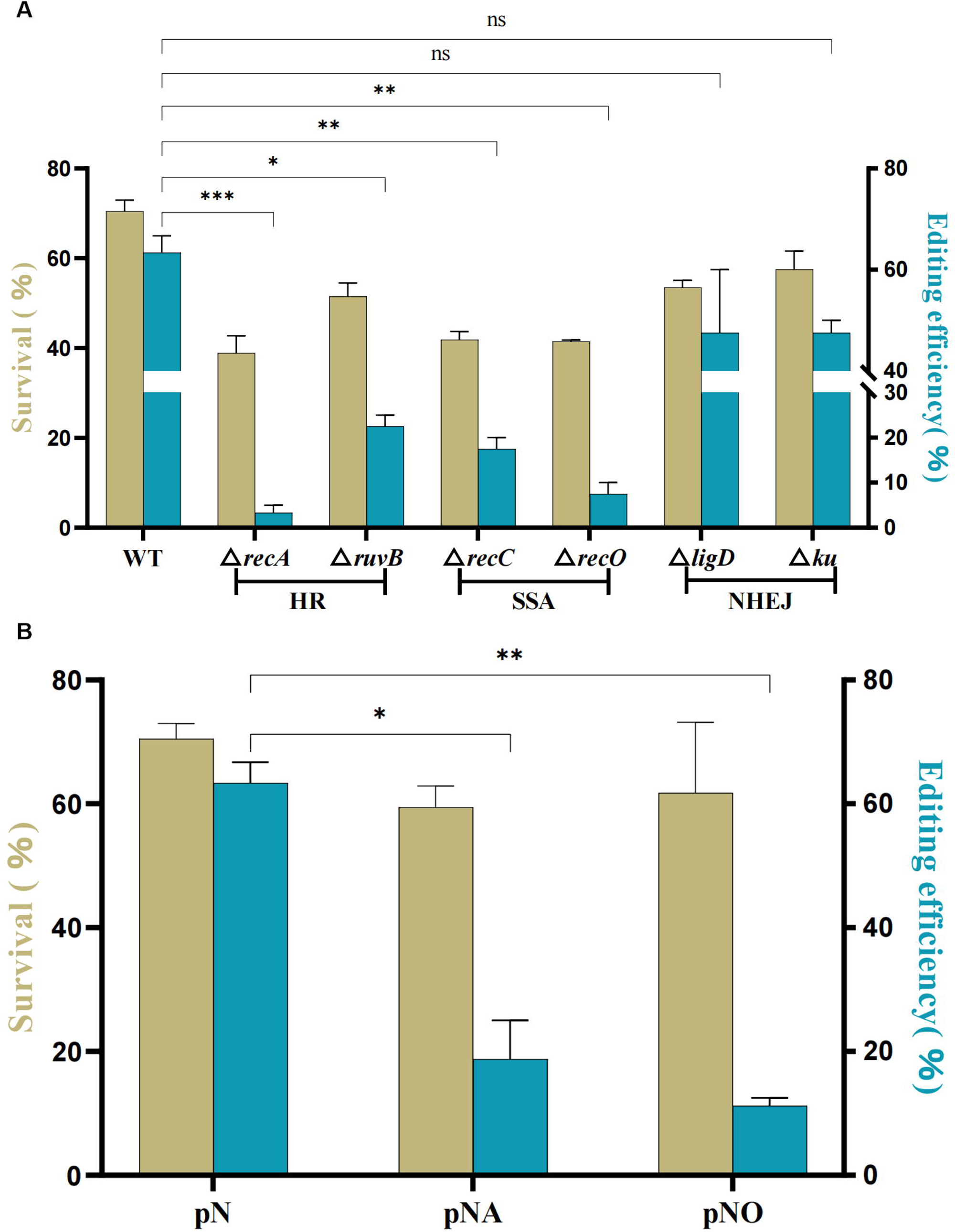
CRISPR-Cas12a-NHEJ gene editing efficiency and survival in *M. abscessus* and its derivatives. Plasmids carrying the crRNA were electroporated into *M. abscessus* and its derivatives, knocking out the key genes in HR or SSA (A), overexpressing *recA* and *recO* gene (B) all resulted in a significant decrease in gene editing efficiency in *M. abscessus*. Editing efficiency was calculated by measuring the proportion of colonies with mutations at the target site among the total randomly selected clones and survival rate was defined as the ratio of the number of CFUs obtained from plates with ATc to the number of CFUs obtained from plates without ATc. ns, not significant. *, *P* < 0.05. **, *P* < 0.01. ***, *P* < 0.001.

In the *ligD* or *ku* gene-knockout strains of *M. abscessus*, the gene editing efficiency compared to the wild-type strains was 47.5 ± 17.68% and 47.5 ± 3.536%, respectively. Although the difference in editing efficiency between these strains and the wild-type strains was insignificant, there was still a slight decrease. This is consistent with our previous results, as we found that when only *nrgA* was introduced, the endogenous NHEJ in *M. abscessus* was capable of handling low-efficiency DSB repair (Fig. 2B). Therefore, when we knocked out the *ligD* or *ku* genes, this part of the repair that was reliant on endogenous NHEJ in *M. abscessus* was eliminated, leading to a slight decrease in gene editing efficiency compared to the wild-type.

Interestingly, editing efficiencies of the knock-out strains ranked as follows: *recA* (HR) ≤ *recO* (HR&SSA) ≤ *recC* (SSA)≤ *ruvB* (HR). A significant difference in editing efficiencies was observed between *recA* and *recC* (*P* = 0.0156), *recA* and *ruvB* (*P* = 0.0067). This suggests that NHEJ-dependent gene editing efficiency is differentially influenced by the different genes in the HR or SSA DSB repair pathways. Genes involved in the initiation of HR repair may have a greater impact on editing efficiency compared to those involved later in the process. Furthermore, genes that participate in HR pathways may exert a stronger influence on editing efficiency than those affecting only SSA. While *recO* might influence both HR and SSA, it primarily affects a branch pathway of HR, unlike *recA*, which plays a central role in HR^28, 30, 31^.

Based on the above results, we found that inhibiting the HR and SSA pathways in *M. abscessus* leads to a significant decrease in NHEJ repair efficiency. This raises the question: if we instead enhance the HR or SSA pathways, could we thereby improve the efficiency of NHEJ repair?

We further overexpressed the *recA* and *recO* genes and repeated the gene editing experiment. Contrary to our expectations, overexpressing either *recA* or *recO* did not enhance the gene editing efficiency based on NHEJ repair. Instead, in *recA* and *recO* overexpressing groups, the gene editing efficiency significantly decreased compared to the wild-type strain, to 18.75 ± 8.839%, 11.25 ± 1.768%, respectively. This suggests that both excessive enhancement of HR or SSA and inhibition of HR or SSA pathways lead to a decrease in NHEJ-based editing efficiency. This decline may be due to the disruption of the interplay among HR, SSA, and NHEJ repair pathways. In summary, HR, SSA, and NHEJ are likely not simply competing pathways but rather interact with each other during the repair of DSBs, existing in a dynamic equilibrium.

## DISCUSSION

*M. abscessus*, a prevalent NTM, has been responsible for a rising number of infections in recent years, attracting widespread concern. More worryingly, *M. abscessus* exhibits intrinsic resistance to many commonly used drugs, leading to a decrease in the effectiveness of current treatment regimens, which makes treating *M. abscessus*-induced infections extremely difficult and poses a serious threat to public health^5, 32^. The issue of drug resistance challenges existing antimicrobial treatment strategies, but on the other hand, it also prompts the scientific community to accelerate research and development of new treatment methods. A deep understanding of *M. abscessus*’s mechanisms of resistance and drug targets will provide a scientific basis for designing more powerful drugs and effective treatment regimens. Therefore, developing specialized genetic manipulation tools to rapidly and efficiently obtain *M. abscessus* gene knockout strains is crucial^33,34^.

In our initial studies, we successfully achieved gene editing in *M. abscessus* by enhancing the recombination frequency through the expression of the *gp60* and *gp61* from the Mycobacteriophage Che9c, or by using the CRISPR-Assisted recombineering method^19, 20^. However, these methods have their obvious limitations: (1). Recombineering-based gene editing methods are relatively inefficient. Our primary results indicate that among the colonies selected on the plates obtained through these methods, those that have undergone gene editing sometimes account for less than 10% of the total selected colonies, which significantly increases our workload in screening for gene-edited strains. (2) Methods based on recombineering require the involvement of an additional homologous template. Gene editing mediated by HR needs a complete template to guide the repair process, necessitating the preparation of additional homologous templates. (3) Mutants obtained by recombinant methods without CRISPR-Cas system will leave an additional resistance gene in the genome and will remain scarred after deletion of the resistance gene^16^. The limitations of these methods make it difficult to efficiently obtain large numbers of *M. abscessus* knockout mutants in a short period of time.

NHEJ, as a DSB repair mechanism that is not common in bacteria, demonstrates great potential for genetic manipulation in combination with the CRISPR-Cas system due to its template-independent and error-prone repair characteristics. However, the mere introduction of the CRISPR-Cpf1 system does not directly enable gene editing based on NHEJ repair in *M. abscessus*, even though *M. abscessus* itself possesses an NHEJ system. Previously, gene editing based on NHEJ repair was achieved in *M. marinum* by solely expressing the CRISPR-Cas system, proving that the NHEJ in *M. marinum* might be a potent repair system^18^. Compared to the NHEJ repair system in *M. abscessus*, the NHEJ components in *M. marinum* contain an additional *nrgA* gene, this may be one of the reasons why *M. marinum* can achieve gene editing based on its endogenous NHEJ repair by solely expressing the CRISPR-Cas system. Therefore, we attempt to combine the CRISPR-Cas system and *M. marinum* NHEJ repair to achieve efficient gene editing in *M. abscessus*. The experimental results of using this method to knock out the *MAB_3513c* gene indicate that the method is highly efficient, with a gene editing efficiency exceeding 60%. The obtained mutants all experienced deletions of base pairs at the target site, but the length of deletion varied, allowing for the selection of strains with different types of mutations according to actual needs. Furthermore, we have further knocked out more than 30 *M. abscessus* genes using this method, with an overall gene editing efficiency ranging between 40% and 90%. This is sufficient to meet the needs for obtaining *M. abscessus* mutants and has significantly accelerated the progress of our research. In any case, we have successfully achieved efficient gene editing in *M. abscessus* for the first time by combining the CRISPR-Cas system with the *M. marinum* NHEJ repair mechanism.

Compared to the NHEJ in *M. marinum*, the NHEJ components in *M. abscessus* only include LigD and Ku, but not NrgA. Previous studies have shown that the presence of NrgA enhances NHEJ repair efficiency in both *M. marinum* and *M. tuberculosis*, leading us to believe that NrgA might play a critical role in the repair of CRISPR-Cas-induced DSBs in *M. abscessus* through NHEJ^18^. A series of experiments expressing only partial NHEJ components from *M. marinum* on plasmids confirmed our hypothesis. NrgA appears to synergize with NHEJ repair in *M. marinum* in some manner. Interestingly, NrgA can also collaborate with the endogenous NHEJ repair in *M. abscessus*, enabling low-efficiency gene editing.

Furthermore, we also discovered that the compatibility of NHEJ components could be a key factor affecting NHEJ repair efficiency. Whether we mixed *ligD* and *ku* from *M. abscessus* or *M. marinum*, or paired *ligD* and *ku* from *M. abscessus* with *nrgA* from *M. marinum*, the gene editing efficiency significantly decreased compared to expressing the complete NHEJ components from *M. marinum*.

NrgA, as a type of phosphofructokinase B (PfkB), has had its structure resolved, making it an intriguing question whether it regulates NHEJ repair by altering cellular metabolism^22^. A study on *M. tuberculosis* pointed out that PfkB is upregulated under survival pressures such as hypoxic conditions and is not negatively regulated by high levels of ADP and ATP^35^. As for NrgA, whether it regulates energy metabolism to produce more dNTPs or rNTPs to supply substrates for NHEJ repair remains to be further studied.

Previous studies largely indicated that the three DSB repair mechanisms compete against each other for gene editing in mycobacteria^18, 25^, which contrasts sharply with our findings in this study. First, a study showing that disruption of HR would not increase the NHEJ-mediated DSB repair efficiency in *M. smegmatis* supports what we found here that disruption of HR did not affect the CRISPR-Cas12a or CRISPR-Cas9 assisted NHEJ gene editing efficiency in *M. smegmatis*^13^ (Fig. 1A). And for *M. abscessus,* our primary results indicate that the CRISPR-NHEJ strategy can still be successfully applied for efficient gene editing in *M. abscessus* without inhibiting the HR pathway. With further research, we found that, regardless of whether in *M. abscessus*’s HR or SSA repair-deficient strains, the efficiency of gene editing based on NHEJ repair significantly decreased, revealing a potential synergistic interaction between HR or SSA and NHEJ within this bacterium. This is consistent with recent research findings in *Halomonas bluephagenesis*, which also indicate that inhibiting HR will significantly reduce the gene editing efficiency based on NHEJ repair^36^. After overexpressing the key genes of the HR or SSA pathway, namely *recA* or *recO*, in *M. abscessus*, gene editing efficiency also significantly decreased. This implies that if the HR or SSA pathway is overly active, it can similarly inhibit NHEJ repair. Based on these results above, we believe that the three types of DSB repair mechanisms in *M. abscessus* might exist in a dynamic balance: similar to a three-legged stool, only when HR, SSA, and NHEJ are in a relatively balanced state can DSBs be effectively repaired. Conversely, take HR as an example, when it is either too weak or too strong, this balance will be disrupted, ultimately leading to inefficient DSB repair (Fig 5). In the future, it might be possible to use promoters of varying strengths to enhance HR or SSA, in order to determine under what strength the NHEJ repair system is inhibited or enhanced.

**FIG 5.**
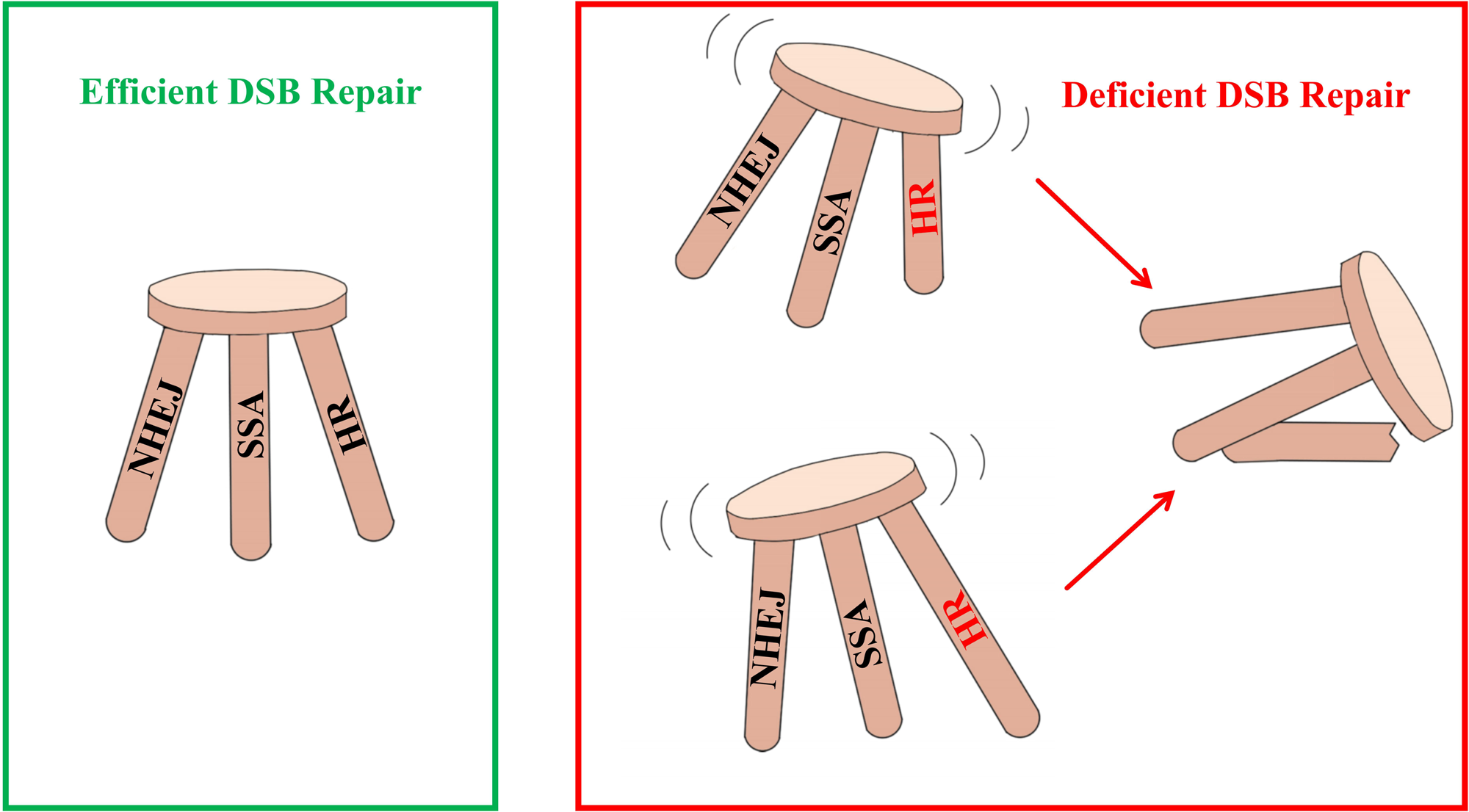
A cartoon representation of the hypothesis concerning the relationship between the three DSB repair pathways. Similar to a three-legged stool, the effective repair of DSBs occurs only when HR, SSA, and NHEJ are in a relatively balanced state. On the other hand, using HR as an example, if it becomes either too weak or too strong, this balance will be disrupted, leading to inefficient DSB repair.

It is interesting to consider whether the differences observed are related to distinct evolutionary branches, which is particularly supported by a recent study reclassified *M. abscessus* and *M. smegmatis* into different new genera, namely Mycobacteroides and Mycolicibacterium, respectively^37^. Our novel finding offers a new perspective for further exploration of the interrelationships between DSB repair mechanisms in mycobacteria and paves the way for future enhancement of specific repair pathway efficiencies by manipulating these relationships.

It is noteworthy that different CRISPR effector proteins produce DSBs of varying forms. For example, the two commonly used Class 2 CRISPR effector proteins, Cpf1 and Cas9, generate different types of DSBs. Cpf1 generates a DSB with a 5′ overhang, in contrast to the blunt ends generated by Cas9^7^. The pathway and outcome of NHEJ repair of DSBs depend on the structure of the broken DNA ends^38^. The processing of different types of DNA ends may involve complex interactions among HR, SSA, and NHEJ. Moreover, different DSB repair preferences for various forms of broken DNA ends may also vary^13^. All these factors could potentially be reasons that affect the relationships among the three DSB repair systems. More refined research on the relationship among HR, SSA and NHEJ, and their impact on the CRISPR-assisted NHEJ gene editing efficiency in different mycobacteria requires further investigation.

## MATERIALS AND METHODS

### Strains, media, and growth conditions

The *E. coli* Tran1-T1 strain was used for molecular cloning and plasmid manipulation. Mab subsp. abscessus GZ002^39, 40^ used in this experiment originated from Guangzhou Chest Hospital. *M. abscessus* derived strains and *M. smegmatis* MC^2^ 155 were preserved by our laboratory. The Luria-Bertani (LB) medium (10 g/L tryptone, 5 g/L yeast extract, 10 g/L NaCl) was used for the culture of *E. coli*. The 7H9 medium supplemented with 10% oleic acid-albumin-dextrose-catalase (OADC) and 0.05% Tween80 was used for the liquid culture of *M. abscessus* and *M. smegmatis*. The 7H11 medium supplemented with 10% OADC and 0.05% Tween80 was used for the solid culture of *M. abscessus* and *M. smegmatis* after electroporation. The working concentrations of anhydrotetracycline (ATc), kanamycin and zeocin were 200 ng/mL, 100 μg/mL and 30 μg/mL for *M. abscessus*, 50 ng/mL, 50 μg/mL and 30 μg/mL for *M. smegmatis*, respectively.

### Construction of plasmids

The primers used for plasmid construction in this study were listed in Table S1. The plasmids related to the CRISPR-Cpf1-NHEJ gene editing system were pCR-Zeo, pNHEJ-Cpf1, and pDK-Cpf1, pK-Cpf1, pA-Cpf1, pD-Cpf1, pAK-Cpf1, pDA-Cpf1, pNHEJ-Cpf1 (*ligD*^Mab^), pNHEJ-Cpf1 (*ku*^Mab^), pNHEJ-Cpf1 (NHEJ^Mab^+*nrgA*^Mmr^), pNHEJ-Cpf1 (NHEJ^Mab^), pNHEJX^Mab^-Cpf1 and pNHEJX^Msm^-Cpf1. The plasmids related to the CRISPR-Cas9-NHEJ gene editing system were pZEO2085, pNHEJ-Cas9 and pNHEJ-Cas9X^Msm^. The plasmid fragments containing *cpf1* originated from the plasmid pJV53-Cpf1. The NHEJ components of *M. abscessus*, *M. marinum*, *recX* from *M. abscessus* and *M. smegmatis* or *recA* and *recO* from *M. abscessus* were amplified using the corresponding primers and ligated into the plasmid backbone containing *cpf1* through recombination methods. The crRNA or sgRNA sequence was incorporated into a pair of primers, which were subsequently annealed and ligated into the linearized pCR-Zeo or pZEO2085 plasmid.

### Gene editing in M. abscessus and M. smegmatis

*M. abscessus* and its derivative strains or *M. smegmatis* carrying the respective plasmids were cultured in shake flasks containing 40 mL of complete 7H9 medium at 37_, 200 rpm until the OD_600_ = 0.8∼1.0. After that, the bacteria were harvested for competent cell preparation as previously described^41^. About 500 ng of the plasmid expressing guide RNA (primers for constructing crRNA are listed in Table S1) was mixed with 100 μL of competent cells, transferred to a 2 mm electroporation cuvette, and incubated on ice for 10 min. Then the cuvette was placed into an electroporation chamber for electroporation at a voltage of 2,500 V, a capacitance of 25 μF, and a resistance of 1,000 Ω. After electroporation, 2 mL of 7H9 medium was added to the electroporation cuvette to resuspend the bacteria, which was then transferred to a 50 mL tube and incubated at 30°C for 4 to 5 hours. Then, cultures were plated separately on 7H11 agar supplemented with the appropriate antibiotics and anhydrotetracycline (ATc) at 30°C. The editing efficiency was determined by the ratio of colonies with altered target sequences to the total number of randomly picked colonies and survival rate was defined as the ratio of the number of CFUs obtained from plates with ATc to the number of CFUs obtained from plates without ATc. For each assessment, at least 20 colonies were randomly selected for PCR (primers are shown in Table S1) and sequencing analysis. Colonies that either lacked the target bands in PCR or showed sequencing results that deviated from the wild-type sequence were identified as having undergone gene editing.

## ACKNOWLEDGMENTS

This study was supported by the National Key R&D Program of China (2021YFA1300900), the National Natural Science Foundation of China (21920102003), the State Key Lab of Respiratory Disease, Guangzhou Institute of Respiratory Diseases, First Affiliated Hospital of Guangzhou Medical University (SKLRD-Z-202301, SKLRD-Z-202412), and Guangzhou Science and Technology (2024A04J4273), the Chinese Academy of Sciences, President’s International Fellowship Initiative (2023VBC0015). The funders had no role in study design, data collection and analysis, decision to publish, or preparation of the manuscript. All authors read and approved the final version of the manuscript.

We also extend our sincere appreciation to Professor Yichen Sun from the Chinese Academy of Medical Sciences and Peking Union Medical College for generously providing the pJV53-Cpf1 and pCR-Zeo plasmids for our study.

## AUTHOR CONTRIBUTIONS

**Sanshan Zeng**: conceptualization (equal); data curation(lead); formal analysis (lead); investigation (lead); methodology(lead); writing — original draft(lead); writing — review and editing (equal). **Yanan Ju**: conceptualization (equal) and investigation (equal); **Md Shah Alam**: conceptualization (equal) and investigation (equal); **Ziwen Lu**: methodology(equal) and software(equal); **H.M. Adnan Hameed**: writing— original draft(supporting) and writing — review(lead); **Lijie Li**: methodology(supporting) and software(supporting); **Xirong Tian**: validation(supporting) and formal analysis(supporting); **Cuiting Fang**: validation(supporting) and formal analysis(supporting); **Xiange Fang**: validation(supporting) and investigation(supporting); **Jie Ding**: validation(supporting) and investigation(supporting); **Xinyue Wang**: validation(supporting) and writing— review(equal); **Jinxing Hu**: validation(supporting) and writing—review(supporting); **Shuai Wang**: project administration(lead) and funding acquisition(lead); **Tianyu Zhang**: project administration(lead) and funding acquisition(lead);

## ETHICS STATEMENT

This study did not involve any human participant or animal subject.

---

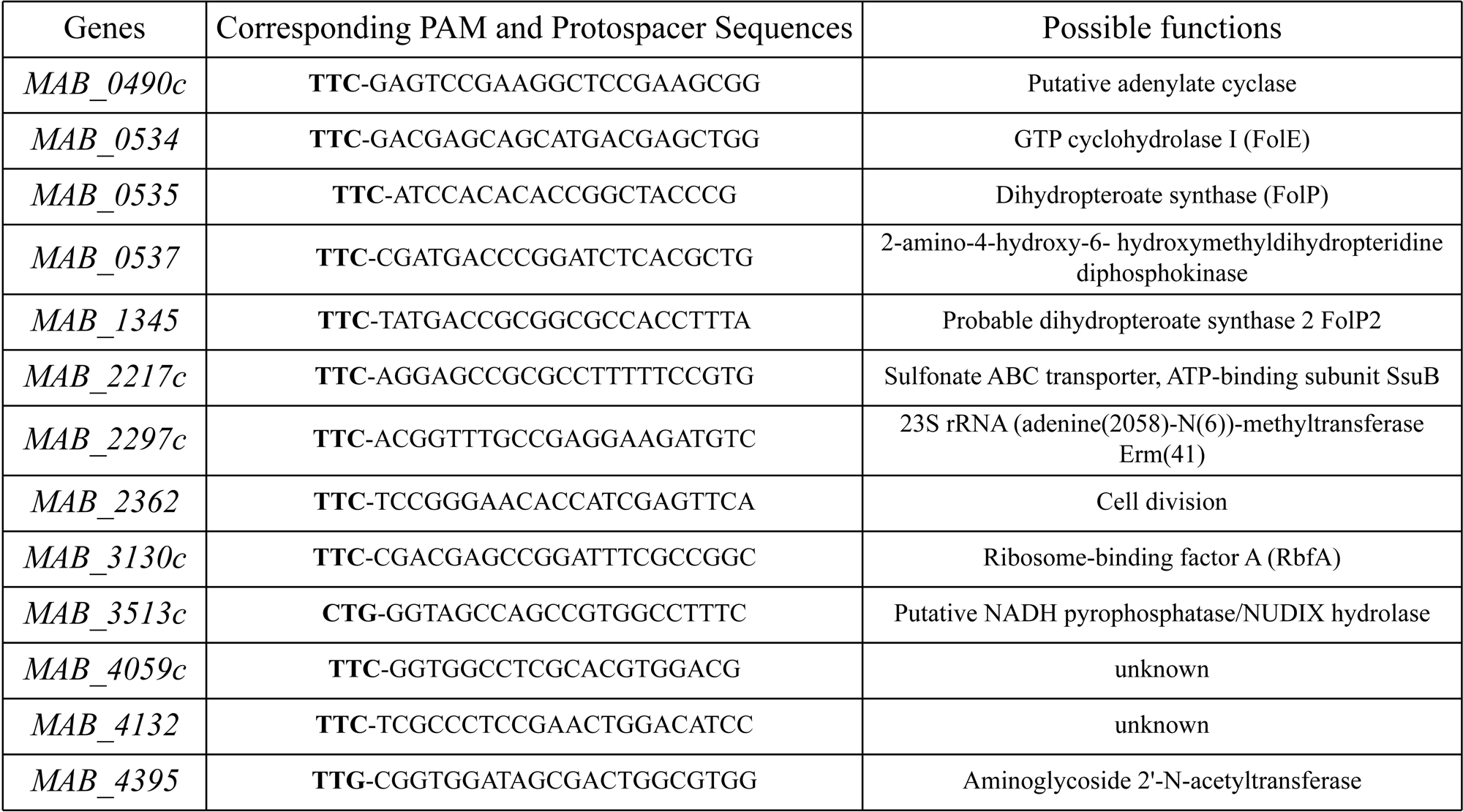

