## Supplementary material for "A CRISPR-Nonhomologous End-Joining-based strategy for rapid and efficient gene disruption in *Mycobacterium abscessus*": FIG S1, Supplementary Table 1 and Supplementary Table 2

**FIG S1.** Maps of plasmid pNHEJ-Cpf1 developed for CRISPR-Cpf1-NHEJ-assisted gene editing in *M abscessus and M smegmatis*. *ligD*, *nrgA*, and *ku* compose the *M marinum* NHEJ element. *kanR*, confers resistance to kanamycin in bacteria. *tetR*, encodes the tetracycline repressor protein, which regulates the expression of gene downstream of the tetracycline operator. *cpf1*, a CRISPR-associated endonuclease used for cleaving specific DNA sequences.

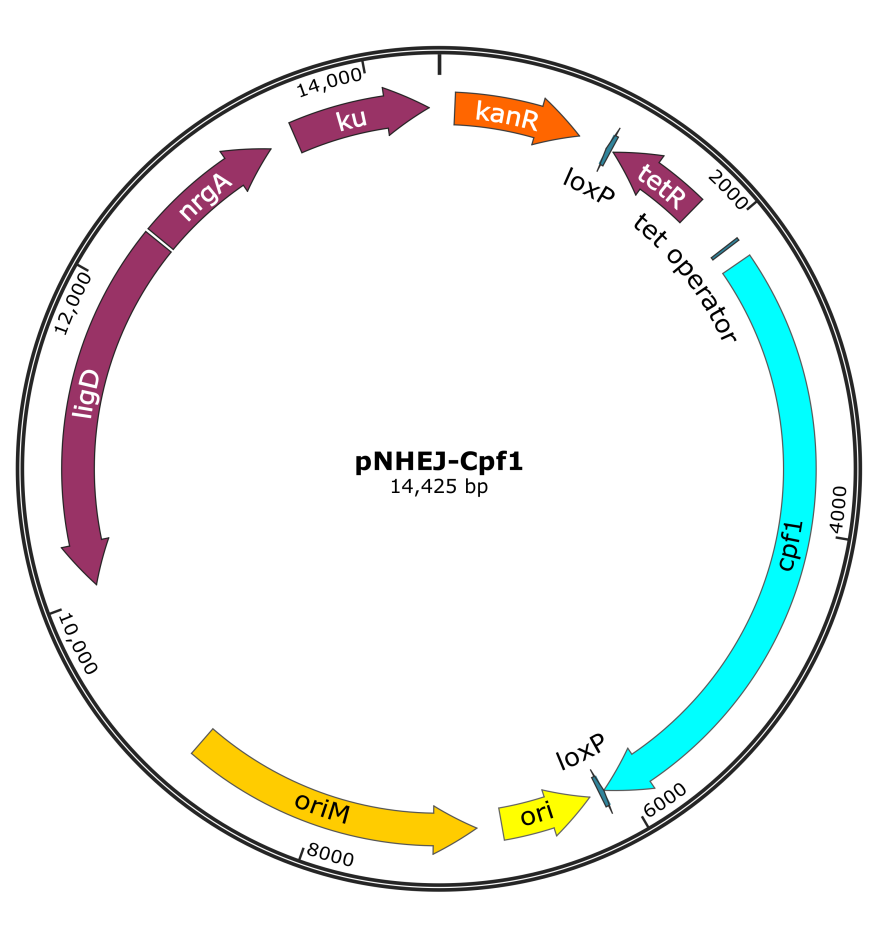

**Supplementary Table 1.** Plasmids used in this study.

| **Plasmid** | **Relevant characteristics** | **Source** |
| --- | --- | --- |
| pJV53-Cpf1 | Used for gene knockout in *M abscessus* and *M smegmatis* based on the CRISPR-Cpf1-HR strategy; Cpf1 is expressed under a tetracycline-inducible system, while gp60/61 is expressed under acetamide induction; Kanamycin resistance. | [1] |
| pNHEJ-Cpf1 | Used for gene knockout in *M abscessus* and *M smegmatis* based on the CRISPR-Cpf1-NHEJ strategy; Cpf1 is expressed under a tetracycline-inducible system, while *M marinum* NHEJ is expressed constitutively; Kanamycin resistance. | This study |
| pDK-Cpf1 | pNHEJ-Cpf1-derived plasmid; Cpf1 is expressed under a tetracycline-inducible system, while *M marinum* *ligD* and *ku* are expressed constitutively; Kanamycin resistance. | This study |
| pK-Cpf1 | pNHEJ-Cpf1-derived plasmid; Cpf1 is expressed under a tetracycline-inducible system, while *M marinum* *ku* is expressed constitutively; Kanamycin resistance. | This study |
| pA-Cpf1 | pNHEJ-Cpf1-derived plasmid; Cpf1 is expressed under a tetracycline-inducible system, while *M marinum* *nrgA* is expressed constitutively; Kanamycin resistance. | This study |
| pD-Cpf1 | pNHEJ-Cpf1-derived plasmid; Cpf1 is expressed under a tetracycline-inducible system, while *M marinum* *ligD* is expressed constitutively; Kanamycin resistance. | This study |
| pDA-Cpf1 | pNHEJ-Cpf1-derived plasmid; Cpf1 is expressed under a tetracycline-inducible system, while *M marinum* *ligD* and *nrgA* are expressed constitutively; Kanamycin resistance. | This study |
| pAK-Cpf1 | pNHEJ-Cpf1-derived plasmid; Cpf1 is expressed under a tetracycline-inducible system, while *M marinum* *nrgA* and *ku* are expressed constitutively; Kanamycin resistance. | This study |
| pNHEJ-Cpf1 (*ligD*^Mab^) | pNHEJ-Cpf1-derived plasmid; Cpf1 is expressed under a tetracycline-inducible system, while *M marinum* *nrgA* and *ku*, *M abscessus ligD* are expressed constitutively; Kanamycin resistance. | This study |
| pNHEJ-Cpf1 (*ku*^Mab^) | pNHEJ-Cpf1-derived plasmid; Cpf1 is expressed under a tetracycline-inducible system, while *M marinum* *nrgA* and *ligD*, *M abscessus ku* are expressed constitutively; Kanamycin resistance. | This study |
| pNHEJ-Cpf1 (NHEJ^Mab^+*nrgA*^Mmr^) | pNHEJ-Cpf1-derived plasmid; Cpf1 is expressed under a tetracycline-inducible system, while *M marinum* *nrgA* and *ligD*, *M abscessus ku* and *ligD* are expressed constitutively; Kanamycin resistance. | This study |
| pNHEJ-Cpf1 (NHEJ^Mab^) | pNHEJ-Cpf1-derived plasmid; Cpf1 is expressed under a tetracycline-inducible system, while *M abscessus ku* and *ligD* are expressed constitutively; Kanamycin resistance. | This study |
| pNHEJX-Cpf1(*recX*^Mab^) | pNHEJ-Cpf1-derived plasmid; Cpf1 is expressed under a tetracycline-inducible system, while *M abscessus recX* is expressed constitutively; Kanamycin resistance. | This study |
| pNHEJX-Cpf1(*recX*^Msm^) | pNHEJ-Cpf1-derived plasmid; Cpf1 is expressed under a tetracycline-inducible system, while *M smegmatis recX* is expressed constitutively; Kanamycin resistance. | This study |
| pNHEJ-SacB | pNHEJ-recX-sacB-derived plasmid; *M marinum* NHEJ is expressed constitutively; Kanamycin resistance. | This study |
| pNHEJ-SacB(*recX*^Msm^) | pNHEJ-recX-sacB-derived plasmid; *M smegmatis recX* is expressed constitutively, while *M marinum* NHEJ is expressed constitutively; Kanamycin resistance. | This study |
| pCR-Zeo(carrying crRNA) | Used for expressing crRNA. Zeocin resistance. | This study |
| pZEO2085(carrying sgRNA) | Used for expressing sgRNA. Zeocin resistance. | This study |

**Supplementary Table 2.** Primary primers used in this study (nucleotides **in bold** indicate genomic sequences, the remainder represent the recombinant sequences).

| Primers | Sequences (all 5’ to 3’) | Note |
| --- | --- | --- |
| MmNHEJ-F | GTAGTCTGCGCTTCAAAGCTT**ACAACACCCCGACACGCTCAC** | The forward primer amplifies the *M marinum* NHEJ element for the construction of the pNHEJ-Cpf1 or pNHEJX-Cpf1 plasmids. |
| MmNHEJ-R | TGAGACACAA**GACTTCACGCAAAAACTCCTCTC** | The reverse primer amplifies the *M marinum* NHEJ element for the construction of the pNHEJ-Cpf1 or pNHEJX-Cpf1 plasmids. |
| Kan-F | GCGTGAAGTC**TTGTGTCTCAAAATCTCTGATGTTACA** | The forward primer amplifies *kanR* fragment for the construction of the pNHEJ-Cpf1 or pNHEJX-Cpf1 plasmids. |
| Kan-R | ATTATACGAAGTTATGAATTC**TTGTAGGTGGACCAGTTGGTGA** | The reverse primer amplifies *kanR* fragment for the construction of the pNHEJ-Cpf1 or pNHEJX-Cpf1 plasmids. |
| HSP-F | CCGTGGCGCGGCCGCGGTACC**GGTGACCACAACGACGCG** | The forward primer amplifies *hsp* promoter for the construction of the pNHEJX-Cpf1 plasmid. |
| HSP-R(Mab-recX) | AGGACGTCAT**CGCAATTGTCTTGGCCATTG** | The reverse primer amplifies *hsp* promoter for the construction of the pNHEJX-Cpf1(*recX*^Mab^) plasmid. |
| HSP-R(Mab-recA) | TGCGCCAT**CGCAATTGTCTTGGCCATTG** | The reverse primer amplifies *hsp* promoter for the construction of the pNHEJA-Cpf1 plasmid. |
| HSP-R(Mab-recO) | AAAGCCGCAT**CGCAATTGTCTTGGCCATTG** | The reverse primer amplifies *hsp* promoter for the construction of the pNHEJO-Cpf1 plasmid. |
| HSP-R(Msm-recX) | CGACTTCGTCAT**CGCAATTGTCTTGGCCATTG** | The reverse primer amplifies *hsp* promoter for the construction of the pNHEJX-Cpf1(*recX*^Msm^) plasmid. |
| Mab-recX-F | CAATTGCG**ATGACGAAGTCGTCCCGGC** | The forward primer amplifies *recX*^Mab^ for the construction of the pNHEJX-Cpf1(*recX*^Mab^) plasmid. |
| Mab-recX-R | TAGATTTAAAGATCTGGTACC**TCATCCGACGTCGCGTCG** | The reverse primer amplifies *recX*^Mab^ for the construction of the pNHEJX-Cpf1(*recX*^Mab^) plasmid. |
| Mab-recA-F | AAGACAATTGCG**ATGGCGCAGGCACCGGAT** | The forward primer amplifies *recA*^Mab^ for the construction of the pNHEJA-Cpf1 plasmid. |
| Mab-recA-R | TAGATTTAAAGATCTGGTACC**GGTTCACGCGTCGGGGCT** | The reverse primer amplifies *recA*^Mab^ for the construction of the pNHEJA-Cpf1 plasmid. |
| Mab-recO-F | GACAATTGCG**ATGCGGCTTTATCGGGATCG** | The forward primer amplifies *recO*^Mab^ for the construction of the pNHEJO-Cpf1 plasmid. |
| Mab-recO-R | TAGATTTAAAGATCTGGTACC**GGGAAAACAGCAGGAAACACC** | The reverse primer amplifies *recO*^Mab^ for the construction of the pNHEJO-Cpf1 plasmid. |
| Msm-recX-F | GACAATTGCG**ATGACGTCCTCCCGGCCC** | The forward primer amplifies *recX*^Msm^ for the construction of the pNHEJX-Cpf1(*recX*^Msm^) plasmid. |
| Msm-recX-R | TAGATTTAAAGATCTGGTACC**CTAGACCCGCCGGCGCTC** | The reverse primer amplifies *recX*^Msm^ for the construction of the pNHEJX-Cpf1(*recX*^Msm^) plasmid. |
| MmNHEJ-F(SacB) | GGTGGCATCCGTGGCGCGGCCGC**ACAACACCCCGACACGCTC** | The forward primer amplifies the *M marinum* NHEJ element for the construction of the pNHEJ-SacB plasmids. |
| MmNHEJ-R(SacB) | CGGCGGCACGACGAGCATATG**TGAACAAGTCGGGCGCTG** | The reverse primer amplifies the *M marinum* NHEJ element for the construction of the pNHEJ-SacB or pNHEJX-SacB(*recX*^Msm^) plasmids. |
| MmNHEJ-F2(SacB) | TCAAAGCTT**ACAACACCCCGACACGCTC** | The forward primer amplifies the *M marinum* NHEJ element for the construction of the pNHEJX-SacB(*recX*^Msm^) plasmids. |
| RecX-F(SacB) | GGTGGCATCCGTGGCGCGGCCGC**GGTGACCACAACGACGCG** | The forward primer amplifies *recX*^Msm^ with *hsp* promoter for the construction of the pNHEJX-SacB(*recX*^Msm^) plasmid. |
| RecX-R(SacB) | CGGGGTGTTGT**AAGCTTTGAAGCGCAGACTACAC** | The reverse primer amplifies *recX*^Msm^ with *hsp* promoter for the construction of the pNHEJX-SacB(*recX*^Msm^) plasmid. |
| MAB_3513c-ci-F | AT**GGTAGCCAGCCGTGGCCTTTC**A | Forward primer for constructing crRNA targeting the *MAB_3513c* gene. |
| MAB_3513c-ci-R | AGCTT**GAAAGGCCACGGCTGGCTACC**ATCT | Reverse primer for constructing crRNA targeting the *MAB_3513c* gene. |
| MAB_3513-F | **ATGACCTTCCGCCTTCGC** | Forward primer for amplifying *MAB_3513c* gene. |
| MAB_3513-R | **CTACGGTCTTCCCGCAGGC** | Reverse primer for amplifying *MAB_3513c* gene. |
| MSM_1946-ci-F | AT**GAGACATGCGTGCAGCGCGAG**A | Forward primer for constructing crRNA targeting the *MSMEG_1946* gene. |
| MSM_1946-ci-R | AGCTT**CTCGCGCTGCACGCATGTCTC**ATCT | Reverse primer for constructing crRNA targeting the *MSMEG_1946* gene. |
| MSM_1946-F | **TCCGAGCGTGGTGGAGATGA** | Forward primer for amplifying *MSMEG_1946* gene. |
| MSM_1946-R | **CCGTACCCGCAGGCGTAATT** | Reverse primer for amplifying *MSMEG_1946* gene. |
| MAB_0490c-ci-F | AT**GAGTCCGAAGGCTCCGAAGCGG**A | Forward primer for constructing crRNA targeting the *MAB_0490c* gene. |
| MAB_0490c-ci-R | AGCTT**CCGCTTCGGAGCCTTCGGACTC**ATCT | Reverse primer for constructing crRNA targeting the *MAB_0490c* gene. |
| MAB_0534-ci-F | AT**GACGAGCAGCATGACGAGCTGG**A | Forward primer for constructing crRNA targeting the *MAB_0534* gene. |
| MAB_0534-ci-R | AGCTT**CCAGCTCGTCATGCTGCTCGTC**ATCT | Reverse primer for constructing crRNA targeting the *MAB_0534* gene. |
| MAB_0535-ci-F | AT**ATCCACACACCGGCTACCCG**A | Forward primer for constructing crRNA targeting the *MAB_0535* gene. |
| MAB_0535-ci-R | AGCTT**CGGGTAGCCGGTGTGTGGAT**ATCT | Reverse primer for constructing crRNA targeting the *MAB_0535* gene. |
| MAB_0537-ci-F | AT**CGATGACCCGGATCTCACGCTG**A | Forward primer for constructing crRNA targeting the *MAB_0537* gene. |
| MAB_0537-ci-R | AGCTT**CAGCGTGAGATCCGGGTCATCG**ATCT | Reverse primer for constructing crRNA targeting the *MAB_0537* gene. |
| MAB_1345-ci-F | AT**TATGACCGCGGCGCCACCTTTA**A | Forward primer for constructing crRNA targeting the *MAB_1345* gene. |
| MAB_1345-ci-R | AGCTT**TAAAGGTGGCGCCGCGGTCATA**ATCT | Reverse primer for constructing crRNA targeting the *MAB_1345* gene. |
| MAB_2217c-ci-F | AT**AGGAGCCGCGCCTTTTTCCGTG**A | Forward primer for constructing crRNA targeting the *MAB_2217c* gene. |
| MAB_2217c-ci-R | AGCTT**CACGGAAAAAGGCGCGGCTCCT**ATCT | Reverse primer for constructing crRNA targeting the *MAB_2217c* gene. |
| MAB_2297c-ci-F | AT**ACGGTTTGCCGAGGAAGATGTC**A | Forward primer for constructing crRNA targeting the *MAB_2297c* gene. |
| MAB_2297c-ci-R | AGCTT**GACATCTTCCTCGGCAAACCGT**ATCT | Reverse primer for constructing crRNA targeting the *MAB_2297c* gene. |
| MAB_2362c-ci-F | AT**TCCGGGAACACCATCGAGTTCA** | Forward primer for constructing crRNA targeting the *MAB_2362* gene. |
| MAB_2362c-ci-R | AGCTT**TGAACTCGATGGTGTTCCCGGA** | Reverse primer for constructing crRNA targeting the *MAB_2362* gene. |
| MAB_3130c-ci-F | AT**CGACGAGCCGGATTTCGCCGGC**A | Forward primer for constructing crRNA targeting the *MAB_3130c* gene. |
| MAB_3130c-ci-R | AGCTT**GCCGGCGAAATCCGGCTCGTCG**ATCT | Reverse primer for constructing crRNA targeting the *MAB_3130c* gene. |
| MAB_4059c-ci-F | AT**GGTGGCCTCGCACGTGGACG**A | Forward primer for constructing crRNA targeting the *MAB_4059c* gene. |
| MAB_4059c-ci-R | AGCTT**CGTCCACGTGCGAGGCCACC**ATCT | Reverse primer for constructing crRNA targeting the *MAB_4059c* gene. |
| MAB_4132-ci-F | AT**TCGCCCTCCGAACTGGACATCC**A | Forward primer for constructing crRNA targeting the *MAB_4132* gene. |
| MAB_4132-ci-R | AGCTT**GGATGTCCAGTTCGGAGGGCGA**ATCT | Reverse primer for constructing crRNA targeting the *MAB_4132* gene. |
| MAB_4395-ci-F | AT**CGGTGGATAGCGACTGGCGTGG**A | Forward primer for constructing crRNA targeting the *MAB_4395* gene. |
| MAB_4395-ci-R | AGCTT**CCACGCCAGTCGCTATCCACCG**ATCT | Reverse primer for constructing crRNA targeting the *MAB_4395* gene. |
| MAB_0490c-F | **GTGACTGGCGCGGAGGGT** | Forward primer for amplifying *MAB_0490c* gene. |
| MAB_0490c-R | **TTAGCCGTGACGCGGGCG** | Reverse primer for amplifying *MAB_0490c* gene. |
| MAB_0534-F | **ATGAACTCGCCCGAGACG** | Forward primer for amplifying *MAB_0534* gene. |
| MAB_0534-R | **TCATGTCCTCATCATCAAGTCAAGAACCTCGG** | Reverse primer for amplifying *MAB_0534* gene. |
| MAB_0535-F | **ATGATCGCTGGCGTGACG** | Forward primer for amplifying *MAB_0535* gene. |
| MAB_0535-R | **TCATTTCGGTTCACCTTTGGCGCC** | Reverse primer for amplifying *MAB_0535* gene. |
| MAB_0537-F | **GTGCGTGCCGTGCTTTCGA** | Forward primer for amplifying *MAB_0537* gene. |
| MAB_0537-R | **CTACGTCACCGGACCGG** | Reverse primer for amplifying *MAB_0537* gene. |
| MAB_1345-F | **ATGGCGAGGCGGGCTG** | Forward primer for amplifying *MAB_1345* gene. |
| MAB_1345-R | **TCATGCCAACCCCCTCACTGTTCGC** | Reverse primer for amplifying *MAB_1345* gene. |
| MAB_2217c-F | **ATGACAGCGGTCTTCGAGGTC** | Forward primer for amplifying *MAB_2217* gene. |
| MAB_2217c-R | **AAACACCCAACTGTGCAAGG** | Reverse primer for amplifying *MAB_2217* gene. |
| MAB_2297c-F | **GTGTCCGGCCAACGGTCG** | Forward primer for amplifying *MAB_2297c* gene. |
| MAB_2297c-R | **CAGCGCCGCCTGATCAC** | Reverse primer for amplifying *MAB_2297c* gene. |
| MAB_2362-F | **ATGATCACCCCTATGAACTTGAC** | Forward primer for amplifying *MAB_2362* gene. |
| MAB_2362-R | **TCAGCTGACCAGGTTCTGCAC** | Reverse primer for amplifying *MAB_2362* gene. |
| MAB_3130c-F | **GTGGCAGATCCCGCCCGC** | Forward primer for amplifying *MAB_3130c* gene. |
| MAB_3130c-R | **TCAATCGGGACGGCGCTC** | Reverse primer for amplifying *MAB_3130c* gene. |
| MAB_4059c-F | **ATGACCGCACCAGTTCGCC** | Forward primer for amplifying *MAB_4059c* gene. |
| MAB_4059c-R | **CTAAGCCAGCGCGGAGGC** | Reverse primer for amplifying *MAB_4059c* gene. |
| MAB_4132-F | **ATGACAAACAACCTATTCGTCGG** | Forward primer for amplifying *MAB_4132* gene. |
| MAB_4132-R | **CTAGGCGGCGTGAGCGTC** | Reverse primer for amplifying *MAB_4132* gene. |
| MAB_4395-F | **ATGTCGGCTGTGTCCAATATGC** | Forward primer for amplifying *MAB_4395* gene. |
| MAB_4395-R | **TCACCAGCCGTCGCCGGC** | Reverse primer for amplifying *MAB_4395* gene. |
